# Orthologous marker groups reveal broad cell identity conservation across plant single-cell transcriptomes

**DOI:** 10.1101/2023.06.18.545471

**Authors:** Tran N. Chau, Prakash Raj Timilsena, Sai Pavan Bathala, Sanchari Kundu, Bastiaan O. R. Bargmann, Song Li

**Author notes:** Corresponding authors Correspondence to Tran Chau and Song Li.

## Abstract

Single-cell RNA sequencing (scRNA-seq) technology has been widely used in characterizing various cell types from in plant growth and development^1–6^. Applications of this technology in Arabidopsis have benefited from the extensive knowledge of cell-type identity markers^7,8^. Contrastingly, accurate labeling of cell types in other plant species remains a challenge due to the scarcity of known marker genes^9^. Various approaches have been explored to address this issue; however, studies have found many closest orthologs of cell-type identity marker genes in Arabidopsis do not exhibit the same cell-type identity across diverse plant species^10,11^. To address this challenge, we have developed a novel computational strategy called Orthologous Marker Gene Groups (OMGs). We demonstrated that using OMGs as a unit to determine cell type identity enables assignment of cell types by comparing 15 distantly related species. Our analysis revealed 14 dominant clusters with substantial conservation in shared cell-type markers across monocots and dicots.

## Introduction

To annotate cell clusters of single-cell expression data from diverse plant species, many approaches have been reported, such as using literature-derived marker gene sets^10^, direct integration of data from diverse species^11^, or comparing the gene expression profiles of single-copy orthologous genes between closely related species^12^. However, many cell-type identity marker genes in Arabidopsis are not cell-type specifically expressed in other species^1^. In non-plant systems, a commonly used approach for cell type identification is to integrate single-cell data across species using one-to-one orthologous genes^13^. However, we have found direct integration of single-cell data from diverse plant species resulted in clusters with mixed cell identities (Supplementary Fig. 1, Table S1). Because one-to-one orthologous genes were assigned solely based on sequence similarity, many such genes may have diverged in their cell type-specific expression levels in different plant species^14^.

To address this challenge, we have developed a novel computational strategy called Orthologous Marker Gene Groups (OMGs), which include not only one-to-one but also one-to-many and many-to-many orthologous genes when comparing cell markers across species. Cell type assignments were based on counting overlapping OMGs between a reference single-cell map and a query species without cell cluster annotation. To accurately quantify similarities between cell clusters as well as account for observed marker overlaps that are due to random noise, a statistical test was performed. This test not only relied on the overlapping number of OMGs between two clusters but also considered the total number of non-overlapping OMGs in all other clusters between different species. Because our method does not require data integration, using OMGs to assign cell types to single-cell clusters is fast and efficient. The steps of using the OMG pipeline for cell type annotation are illustrated in Fig. 1a.

**Fig. 1:**
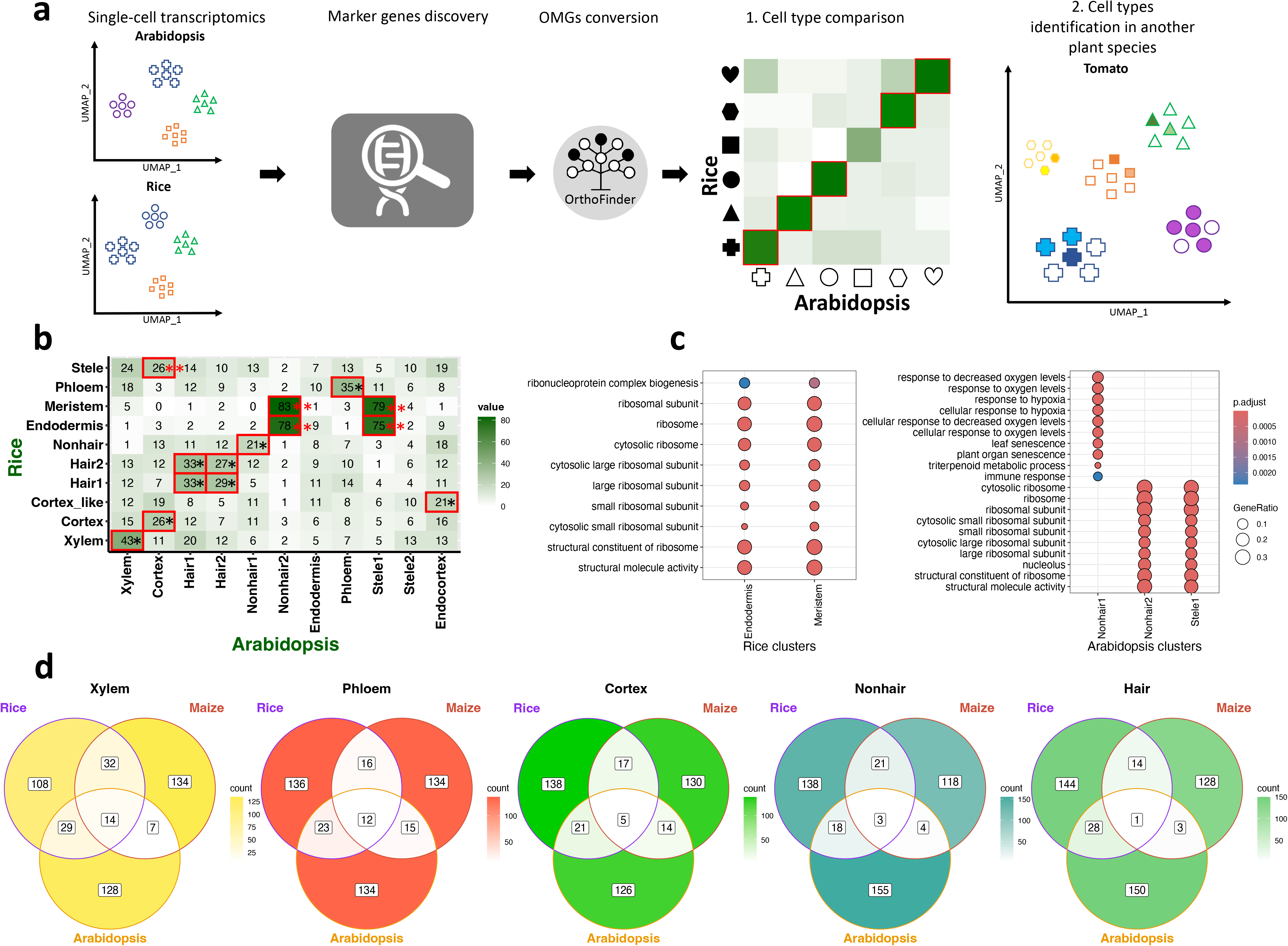
Common OMGs detection. **a**, the OMG pipeline. First, we identified the top N marker genes (N=200) for each cell cluster in each species using an existing approach (e.g. Seurat^21,60^). Second, we employed OrthoFinder^61^ to generate orthologous gene groups for M plant species (M=15, including Arabidopsis, rice (*Oryza sativa*), maize (*Zea mays*), and tomato (*Solanum lycopersicum*), where reference grade single-cell maps are available, and 11 other species where single-cell data have more recently become available). Third, we performed a pairwise comparison using an overlapping OMG heatmap followed by a statistical test (i.e., Fisher’s exact test) to determine clusters with a significant number of shared OMGs. Using N=200 markers per cluster enables consistent statistical comparisons, optimizes overlap of OMGs across species, and maintains marker specificity. Lower N leads to a rapid decrease of overlapping (Supplementary Fig. 5a), and higher N diminishes specificity (Supplementary Fig. 5b). **b**, Pairwise comparison of cell clusters from rice and Arabidopsis. Each box displays the number of conserved OMGs between the two cell types being compared. The red highlighted boxes indicate FDR-adjusted p<0.01 based on Fisher’s exact test. The 9 matched clusters are marked by a black stat (^*^). The putative meristematic clusters are marked by two red stats (^**^). The specificity of the OMG method is illustrated by this heatmap. For example, the Arabidopsis xylem cluster has 43, 14, and 24 OMGs shared with xylem, cortex, and stele clusters in rice. The only significant overlap is between xylem clusters in Arabidopsis and rice. **c**, Gene function analysis of marker genes in Arabidopsis (stele, nonhair) and rice (endodermis, meristem) clusters. The size of the dot indicates the number of genes associated with a particular GO term in the dataset, and the color represents the level of significance of the enrichment. We found genes from the nonhair-1 and nonhair-2 clusters in Arabidopsis do not have the same enriched GO categories. **d**, Conserved OMGs across three species.

To test the power of the OMG method, we used single-cell data from rice^2^ and Arabidopsis^1^ as one example, where 11 clusters in Arabidopsis and 10 clusters in rice have corresponding cell types in the counterpart species. We identified 8 statistically significant cell cluster pairs (FDR < 0.01) using one-to-one orthologous genes, and 14 such pairs using the OMG method (Fig. 1b, and Supplementary Fig. 2). When mapping cell clusters with one-to-one orthologous genes, only 3 out of 8 pairs are from orthologous cell types. In contrast, 9 out of 14 pairs are from identical or nearly identical orthologous cell types based on the OMG method. This result shows that 3 times more clusters were assigned by the OMG method as compared to using one-to-one orthologous genes.

We also observed mismatched clusters with the OMG method, and interestingly, four of these mismatched clusters are in the connecting region of multiple clusters of the Arabidopsis UMAP and the rice UMAP (Supplementary Fig. 3), suggesting these are relatively undifferentiated cell clusters. To test this hypothesis, we compared the root single-cell data from two species with a single-cell atlas with 15 plant species (Supplementary Fig. 4). We found these three clusters are also clustered with the meristematic cells from tomato^4^, cassava roots^15^, wild strawberry (*Fragaria vesca*)^16^, field mustard (*Brassica rapa*)^17^, and Madagascar periwinkle (*Catharanthus roseus*) leaves^18^. Furthermore, using the Gene Ontology (GO) functional enrichment analysis^19^, these three mis-matched clusters from the two species are enriched with similar GO categories, such as ribosomal genes, which are a hallmark of meristematic cell identities (Fig. 1c, Table S2). These results suggest that these clusters are better labeled as meristematic cells. With these clusters re-named as meristematic clusters, the OMG method correctly assigned 13 out of 14 cluster pairs between Arabidopsis and rice, which substantially improved the overall mapping correctness between two distantly related species.

To test whether we can identify OMGs that are conserved in more than two species in monocots and dicots, we used root single-cell data from rice^2^, maize^3^, and Arabidopsis^1^ as our models to define common OMGs. Surprisingly, we found only five cell types contain common OMGs, and for these, only a few OMGs are shared among all three species (Fig. 1d). To tackle this scarcity of common OMGs, we considered increasing the marker gene pool (i.e., N > 200), but this may lower marker specificity (Supplementary Fig. S5a and b). Aiming to bolster the number of markers without sacrificing the specificity, we employed two machine learning methods (SHAP-RF and SVM) and we did not find substantially more OMGs in three species (Supplementary Fig. S6a and b). These results suggest using different methods could lead to the discovery of some new markers. However, there is still a lack of conserved marker genes across these three species, suggesting that pair-wise comparison is a preferred method for cross-species comparison.

To further validate the performance of our OMG method in cell type assignment, we applied OMG method to a single-cell dataset from tomato roots^4^. Tomato has several promoter-GFP lines which provide a gold-standard validation for cell-type identity^4^, and such a resource is not available for most other plant species. Without providing the published cell-type identity from tomato^4^, our analysis found 15 clusters in tomato and compared to the Arabidopsis reference map with the OMG method (Fig. 2a). Among the 15 clusters, the published annotation of 12 clusters in tomato exactly matched the corresponding Arabidopsis clusters (Fig. 2a, Table S3). The cortex cluster in tomato is a “partial match,” because it also shared significant OMGs with both cortex and nonhair cells. There are two exodermis clusters, which is a cell type not found in Arabidopsis. We found that the exodermis clusters in tomato have significant overlapping OMGs with endodermis clusters in Arabidopsis. Both endodermis in Arabidopsis and exodermis in tomato contain suberized barriers in their cell walls^4,20^, which could explain this observed commonality between these two types of cell clusters. In summary, all clusters in the tomato root data showed exact match, partial match, or functional match with corresponding Arabidopsis clusters using the OMG method, and genes from OMG groups showed high cell type specificity (Supplementary Fig. 7).

**Fig. 2:**
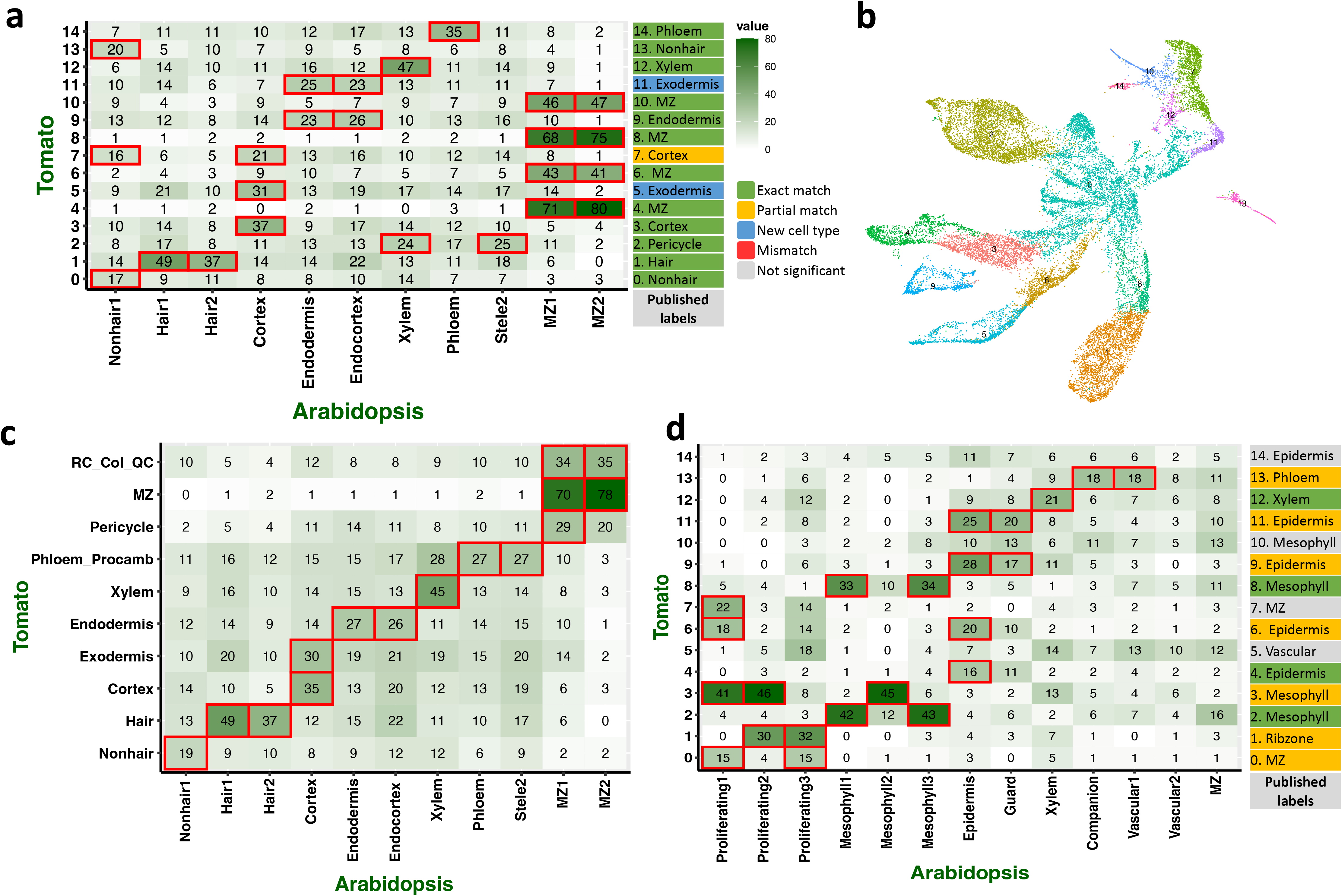
Cell type identification in another plant species. **a**, Using Arabidopsis OMGs to predict tomato root cell types, with the number in each cell reflecting common OMGs between clusters, darker colors indicate more OMGs and red boxes highlight significant overlapping (adjust p value < 0.01). Labels on the right indicate the cell type labels based on published annotation. **b**, UMAP of 15 default tomato root clusters. Specifically, we merged all the meristematic zones (clusters 4, 6, 8, 10, in a) into a single cluster (cluster MZ). Similarly, we have recreated the cortex, hair, and non-hair clusters based on the original assignment in the published study. **c**, pairwise comparison between Arabidopsis and tomato published cell types using published cluster names. **d**, Using Arabidopsis OMGs to predict tomato shoot cell types.

Because the cell clusters were generated using an unsupervised approach, we can adjust the number of clusters based on other criteria. For example, the published UMAP of tomato root cells merged some clusters into bigger clusters of the same cell type (Fig. 2b). To further explore the effect of different cluster numbers, we performed the following tests. First, we adjusted the Seurat^21^ parameter to generate 30 clusters from the tomato dataset (Supplementary Fig. 8a and b). We found that 20 out of 30 clusters are exact matches with the Arabidopsis clusters. An exodermis cluster splits into three clusters with two matches to endodermis and one match to cortex. We also found four clusters without significant matches. This result shows that increasing query cluster numbers maintains a majority of significant overlaps, but also leads to smaller clusters that do not show exact matches to any Arabidopsis clusters, which is due to an insufficient number of marker genes to accurately ascribe cell types. Second, we used the cluster assignment provided by the original publication, and we found ten clusters (Fig. 2b, Table S3). This resulted heatmap also showing high concordance between the cluster annotations in the two species. The same analysis was performed between rice root and tomato root data, and we found fewer significant overlapping OMGs, with the majority of significant similar clusters from the same cell type in two species (Supplementary Fig. 8c).

Expanding the scope of our study to non-root tissues, we tested our method on shoot apical meristem tissues, using Arabidopsis as the reference map to make predictions for tomato. Annotations of these cluster identities were based on the original publications. Eleven out of fifteen clusters showed an exact or partial match in their identity with significant overlap in OMGs (Fig. 2c, Table S3). To further demonstrate the general applicability of this method, we applied this method to single-cell data of roots and shoot apical meristems from rice^2^, maize^3^, poplar^22^, tomato^6^, and Arabidopsis^5^. This concordance of cell types based on OMGs was found between all other pairwise comparisons performed in roots, as well as in shoot apical meristems (Supplementary Fig. 9). These results demonstrate that by using OMGs between different plant species, cell types can be assigned for a majority of cell clusters regardless of the tissue type. The advantage of our method, as compared to those used in published papers^1,4–6^, is that our method is fully automatic and provides a quantitative specificity test. Moreover, our method also allows for fuzzy matches between clusters, in the case of clusters with mixed cell types.

To understand the conservation and divergence of cell identity markers across a variety of plant species, we constructed a comprehensive plant cell atlas that covers 15 species^18^, including Arabidopsis^1,5,8,23–38^, field mustard^17^, Madagascar periwinkle^39^, wild strawberry^40^, soybean^41^, cotton (*Gossypium bickii*^42^ and *Gossypium hirsutum*^43,44^), cassava^45^, barrel clover (*Medicago truncatula*)^46^, coyote tobacco (*Nicotiana attenuata*^47^), rice^24,27,48–50^, *Populus (alba var pyramidalis, and alba x populus glandulosa)*^22,51^, tomato^4,6^, and maize^3,10,52–56^ (Fig. 3a, Table S4). This atlas comprises 268 cell clusters, 990,894 cells, and 53,600 marker genes which were obtained from multiple sources. We clustered cell types from different species based on the shared OMGs across species. One outcome of such mapping would be grouping similar cell types into clusters, neglecting any overriding effects of species dominance. Alternatively, if species-specific effects are significant, cell types from the same species will form species-specific groups. A prior study of mammalian systems demonstrated that gene expression levels group samples by organs whereas in insects, isoform expression levels group samples by species of origin, suggesting a strong species-specific effect on splicing^57,58^.

**Fig. 3:**
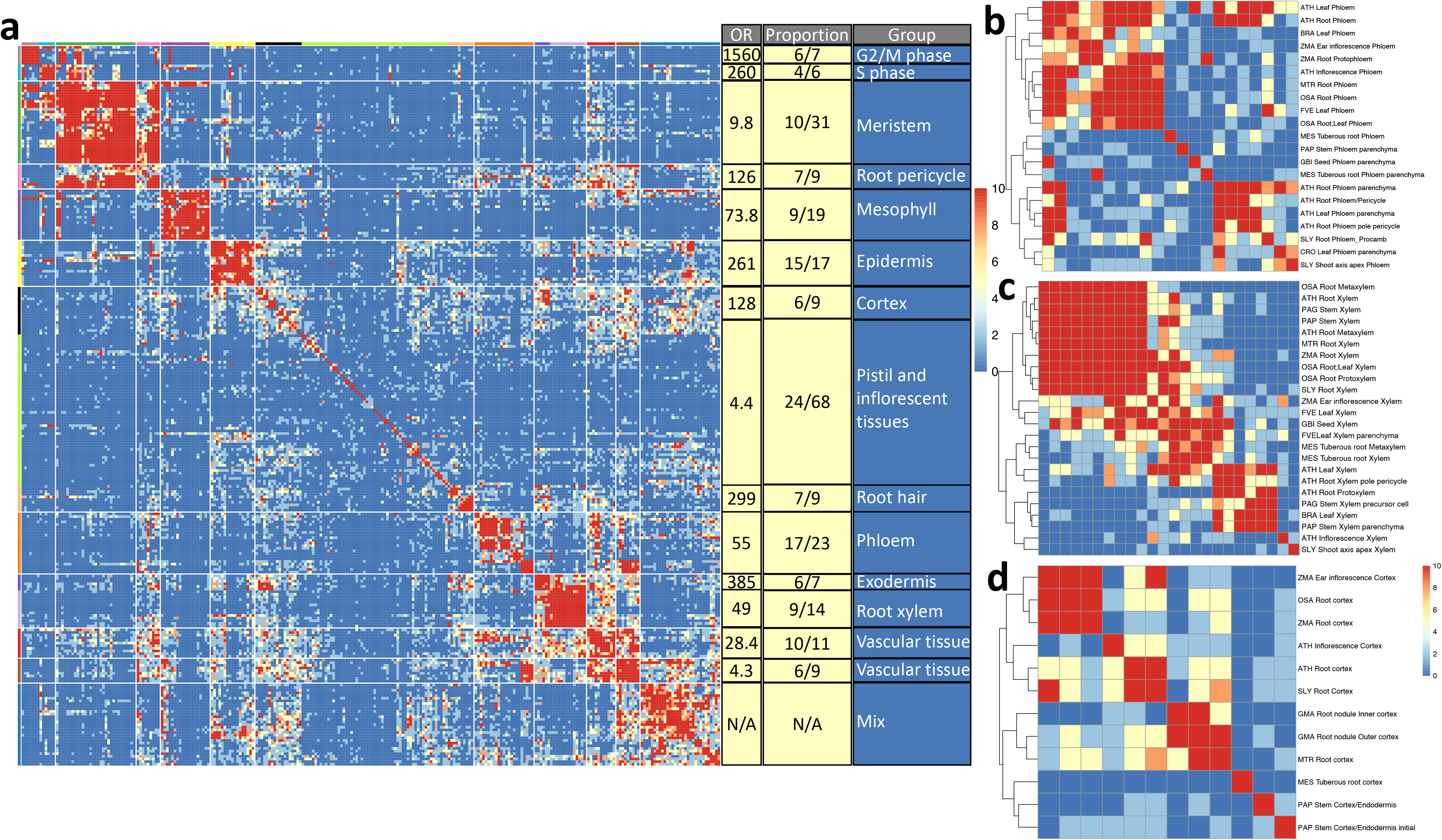
Mapping cell types across 15 diverse species. **a**, pairwise comparison of cell clusters across 15 species based on the shared OMGs. The color scale represents the negative logarithm of the FDR-adjusted p-value. Odd Ratios (OR) represent the likelihood of a particular cell type appearing in a specified group relative to its presence in all other groups. Proportion indicates the frequency of the predominant cell type within each group. Groups are named according to the most prevalent cell type, as indicated by their respective OR and proportion values. **b, c, d**, cell type-specific clustering (phloem, xylem, and cortex) across 15 species.

Our approach identified 15 clusters and 14 of these clusters can be named based on the dominant cell types present within each group as determined by odds ratio (OR range between 4.4 to 1560) and proportion values (Fig. 3a). We observed that all these groups are clustered based on the cell type, not the species. For example, G2M cells from leaf, inflorescence, and shoot apex in different species formed a single cluster. Similarly, the mesophyll cell type is conserved across leaf, shoot apex, and inflorescence in different plant species. Functionally relevant clusters are also closely related in this map, for example, cells from G2M and S phases formed two smaller clusters which are two sub-clusters within a large cluster (Fig. 3a). Detailed annotations of each sub-cluster can be found in Supplementary Fig. 10.

When focused on each cell type, the OMG method revealed intriguing cell type-specific conservation and divergence. For example, the phloem cluster consists of two subclusters, phloem cells and phloem parenchyma/pericycle cells. For phloem cells, there are no separate clusters for leaf phloem or root phloem (Fig. 3b). Interestingly, xylem has a distinct pattern between root and non-root tissues (Fig. 3c), with one sub-cluster consisting of mainly root xylem cells while other sub-cluster consisted of a mixture of xylem cells from different tissue types. The cortex cluster does not show consistent patterns across species, but there is a marked difference between monocots and dicots (Fig. 3d). In the Exodermis cluster, we observed a notable similarity between endodermis in Arabidopsis and exodermis in maize, rice, and tomato, supporting the functional similarity between these cell types (Supplementary Fig. 10). The epidermis in root tissues displays a different pattern compared to non-root tissues: root hair shows a uniform pattern across multiple species, this consistency is not seen in non-hair and root cap cells (Supplementary Fig. 10). Overall, the OMG method offers a robust methodology to map cell types across plant species and reveal tissue-specific patterns. Finally, to understand how the number of common OMGs between two species changes over an evolutionary distance, we found a statistically significant (p<0.05) decrease in common OMGs with increasing evolutionary distance (Supplementary Fig. 11a). However, including species-specific cell types such as cells from soybean nodules eliminate this trend (Supplementary Fig. 11b).

In conclusion, the OMG framework is a straightforward and effective method to attribute cell type identities across multiple plant species using single-cell reference maps. This method is also inherently “explainable” because the lists of overlapping marker genes can be further examined to study their biological relevance (Supplementary Fig. 7). Most cell clusters in our analysis only showed one or two orthologous cell clusters that are statistically significant, suggesting high specificity of this method. However, limitations of this method include that it can only assign cell types to each cluster, and not to individual cells. Different pre-processing methods and the choice of the number of clusters can potentially result in different markers from the same dataset, which creates a hurdle for determining the optimal number of clusters and clustering methods. Finally, there is not enough data to characterize cell type-specific decay of overlapping OMGs (Supplementary Fig.11c). These limitations can be further mitigated with community-based agreement on meta-data publication standards^59^ such that it is easier to reproduce the published clusters, extract marker genes automatically, and enable faster cross-species comparison in many species. The scripts for the analysis can be found at our GitHub repository (https://github.com/LiLabAtVT/OrthoMarkerGenes) and shinyR website for visual comparisons (http://orthomarkergenes.org).

## Online methods

### Cell type assignment for published scRNA-seq datasets

For Fig. 1, gene expression data from *Arabidopsis thaliana, Zea mays*, and *Oryza sativa* were re-analyzed. Parameters for the re-analysis were selected to reproduce UMAP plots that best resemble those in the published results. The data consisted of 7522 Arabidopsis cells, 14,733 *Zea mays* cells, and 27,426 *Oryza sativa* cells, which were published by Ryu et al., Ortiz-Ramírez et al., and Zhang et al., respectively (GSE123013, GSE172302, PRJNA706435). To process the gene expression matrix, a standard workflow that included the NormalizeData, FindVariable, and ScaleData functions, was used with default parameters in the Seurat package (verseion 4.1.1). Principal component analysis (PCA) was used to reduce the high-dimensional data into 30, 15, and 32 principal components (PCs) for Arabidopsis, maize, and rice, respectively. The FindCluster function with resolutions of 0.5, 0.55, and 0.4 was applied to these PCs using the Louvain algorithm to identify the clusters, and the uniform manifold approximation and projection (UMAP) method was used to visualize all cells in two dimensions. Finally, lists of known marker genes from these published papers were used to label all the cell clusters. If a cluster did not include any marker genes from the publication, the cluster is labeled as “unknown”. The cell-type assignment scripts can be found in our GitHub repository.

### Integration of three plant species datasets together

To integrate scRNA-seq datasets from Arabidopsis, maize, and rice, the Seurat integration pipeline was used. First, genes from each species were converted into orthologous gene IDs, and 5000 high variable genes were identified and ranked from 1 to 5000 using the FindVariable function and nfeature=5000. Since one orthologous gene can consist of one or more species of genes, ranking the list of high-variable genes supported the selection of the top genes in each orthologous group for integration. Second, top 2000 (out of 5000) unique high-variable orthologous genes corresponding to the top high variable genes for each species were selected to ensure that the most informative genes were used in the integration process. To ensure the quality of each dataset, the merged Seurat object of three datasets was split into its individual components, and each was independently normalized and scaled using the SCTransform method. The SelectIntegrationFeatures function was then used to identify the top informative features for integrating multiple datasets, and the PreSCTIntegration function was employed to ensure that the Pearson residuals for the selected features were present in each Seurat object. To integrate the three datasets, the FindIntegrationAnchors function was used to identify a set of anchors between all Seurat objects in the list, and these anchors were then used to integrate the three objects together with IntegrateData. Finally, the standard workflow for processing scRNA-seq data was applied, including RunPCA, RunUMAP, FindNeighbors, and FindClusters. The clustering was performed using 30 dimensions and a resolution of 0.3. This analysis produced an integrated UMAP for Arabidopsis, rice, and maize.

The integrated metadata was annotated based on the cell-type annotation obtained from the original analyses where each species was analyzed separately. This step avoided the potential overfitting in the integration process that could lead to loss of biologically important variations across species. To investigate whether the number of homogenous clusters would increase, the resolution parameter of FindClusters was modified to 0.9, resulting in the number of clusters being increased to 30. In addition, to examine the potential impact of heterogenous clusters caused by different integration methods, the harmony integration was employed by applying the RunHarmony function after conducting PCA.

### Marker gene identification by differential gene expression analysis

The FindAllMarkers function in the Seurat package was used to detect marker genes in species-specific UMAPs. A default differential expression test of expression level in a single cell type versus the expression in all other cell types was performed based on the Wilcoxon rank-sum test. To detect marker genes, we included only.pos = TRUE, which selects only positive log-fold changes, min.pct = 0.3, which sets a minimum percentage of cells that express the gene, and logfc.threshold = 0.45, which sets the threshold for log-fold change to select significant genes. After running the FindAllMarkers function, the list of differentially expressed genes was sorted based on their log-fold change values, and the top 200 most significant marker genes were selected for each cluster.

### Marker gene identification by SHAP and SVM methods

We used the SPMarker Python package to identify marker genes in our Arabidopsis, rice, maize, and tomato datasets. The SPMarker package employed two feature selection techniques, namely Shapley additive explanations (SHAP) based on random forest and support vector machine, to develop a pipeline for calculating the contribution of each gene on the cell type assignment model. This approach allowed us to handle imbalanced data caused by major and minor clusters in scRNA-seq data and identify unique marker genes in each cell type cluster. To generate the cell type and gene expression matrix files, a custom R script was used with Seurat’s FindVariable function to select 5000 high-variable genes and the SCTransform function to normalize the selected dataset. Using the SHAP value and SVM coefficients, cell type-specific markers were identified in the species-specific UMAPs. Similar to the default Seurat method, the top 200 most significant marker genes were selected for each cluster.

### Identification of conserved orthologous marker genes and analysis of the conserved orthologous marker genes between two species

OrthoFinder software (v2.5.4) was used to search homologous genes and cluster these genes into ortho-groups. Protein sequences in FASTA format for each species were downloaded from the Ensembl plants database and used as input for OrthoFinder analysis. Species-specific marker gene names were mapped to the ortho-group name and these ortho-groups are named Orthologous Marker gene Groups (OMGs). These OMGs were used in downstream analyses, in place of the original marker genes, to enable comparison between species.

After obtaining lists of orthologous marker genes for all cell types in each of the species, the number of common orthologous marker genes between any two species was calculated. To evaluate the significance of the conserved marker gene list, Fisher’s exact independent tests were performed, hypothesizing that the orthologous marker gene list for each cell-type cluster from one species was independent of those from another species. Fisher’s exact test is a statistical method used to analyze the association between two categorical variables, and in this case, it was used to determine the significance of the overlap in marker genes between species. The p-value from the Fisher test could provide evidence to evaluate the significance of the conserved marker gene list. The p-value is obtained through the R code:

p_value_df[i,j] = fisher.test(rbind(c(a,b),c(c,d)), alternative = “greater”)$p.value

df[i,j] is a matrix where i ∈ (1, M) where M is the number of clusters in species 1 and j ∈ (1, N) where N is the number of clusters in species 2;

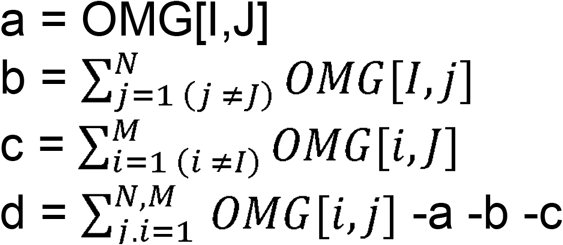

To address the issue of multiple testing, we applied an FDR correction to the p-values obtained from the Fisher exact test. A threshold of 0.01 was then set on the q-value to determine which tests were significant.

### Conservation of OMG across 15 plant species

We employed a hierarchical clustering approach based on Fisher’s exact test p-values to generate a heatmap. The heatmap represents an analysis of a 268 species-specific cell clusters, ∼1 million cells, and over 53,600 marker genes. We began identifying the number of common OMG between cell-type clusters of two species. Data source for the 15 species can be found in Table S5. We then evaluated the significance of the conservation using Fisher’s exact tests, adjusting p-values for FDR to correct for multiple testing. We applied negative log transformation to these p-values at various thresholds for better visual differentiation of significance levels. The “complete” method of hierarchical clustering was applied to define clusters and sub-clusters within the data. The final heatmap was generated by the “pheatmap” R function with color-coded to enhance interpretability. Heatmap was generated using the -log10 FDR adjusted p value rounded to the following thresholds 50, 30, 20, 10, 5, 3, 2 for better visualization.

### Odds ratio (OR) to name cell type groups that map cell types across 15 species

The odds ratio (OR) is a measure indicating the likelihood of a certain cell type across different species clustering together as shown by a heatmap’s color bar, versus scattering among different groups. It is calculated as the ratio of the odds of that cell type appearing in one specific group compared to its odds across all other groups. An OR greater than 1 suggests that the cell type is clustered into a particular group, justifying its use as a label for that group in the heatmap.

### The correlation between phylogenetic distance and shared OMGs between Arabidopsis and other plant species

To investigate this relationship, we processed the phylogenetic tree obtained from OrthoFinder using the “Tree” class in the “ete3” Python module. We normalized the evolutionary distances of each species to Arabidopsis on a scale from 0 to 1, with lower values denoting a closer evolutionary relationship with Arabidopsis and higher values indicating greater divergence. To visualize the relationship between the shared OMGs and the phylogenetic distance, we applied the “ggplot2” package in R to generate a line plot and performed linear regression analysis.

### GO enrichment analysis for Arabidopsis and Rice marker genes

To identify GO terms in our marker gene lists, we performed GO enrichment analysis using the R package such as “clusterProfiler”, “AnnotationDbi”, “AnnotationHub”, along with “org.At.tair.db” for Arabidopsis annotation. We analyzed marker genes associated with Arabidopsis’s nonhair and stele. The “compareCluster” function implemented the enrichment analysis, using a p-value cutoff of 0.01 and adjusted p-value using the Benjamini-Hochberg method. The results were visualized through the “dotplot”, which depicted the gene count by dot size and significance by color. For rice, we accessed the genome annotation via “AnnottaionHub” and used “biomart” to ensure only genes with valid Entrez IDs were included. The enrichment analysis for rice followed the same parameters as for Arabidopsis.

### OMG browser

To facilitate the use of this method by the broad research community, we have launched a user-friendly web-based tool called the OMG browser, which enables effortless identification of cell types or perform pairwise comparison across monocot and dicot. For input preparation, the Seurat package’s FindAllMarker function or alternative methods can be utilized to determine marker genes. Inputs should be formatted as a data frame with three columns specifying gene names, cluster names, and gene expression levels. For guidance on generating this input, please refer to our instructional video at https://youtu.be/oliRmER1rXw. When the marker gene table is uploaded, the browser allows users to conduct pairwise comparisons with our reference data, incorporating a statistical test to highlight significant findings. If the users’ data have not been previously labeled, our browser can assist in labeling through comparison and statistical validation. A detailed tutorial video is available to walk users through the process, which can be found at https://youtu.be/Jb4uMq394Sg.

## Author information

### Authors and Affiliations

**School of Plant and Environmental Sciences, Virginia Tech, Blacksburg, VA, USA** Tran N. Chau, Prakash Raj Timilsena, Sanchari Kundu, Bastiaan O. R. Bargmann & Song Li

**Graduate Program in Genetics, Bioinformatics, and Computational Biology, Virginia Tech, Blacksburg, VA, USA**

Tran N. Chau & Song Li

**Department of Computer Science, Virginia Tech, Blacksburg, VA, USA**

Sai Pavan Bathala & Song Li

## Contributions

S.L. developed the project and supervised its execution. T.N.C. performed the analyses. S.P.B. created the web browser. P.R.T. provided tomato data. S.K. performed the Poplar preprocessing. T.N.C & S.L. wrote the manuscript with contributions from B.O.R.B.

## Peer review

## Supplementary tables

**Table S1:** Seurat integration with 19 clusters, 30 clusters, and Harmony integration.

**Table S2:** Go enrichment analysis of Arabidopsis and rice marker genes. Pairwise comparison of rice vs Arabidopsis using OMG test. In table S2C, the predicted cell types are labeled as “reject” which is the statistical term signify the observed overlapping is not due to random chance, thus the null hypothesis is “rejected”. Other comparisons are listed as “fail”, which means the tests failed to reject the null hypothesis.

**Table S3:** Cell type predictions in tomato root and shoot clusters using OMG tests. Pairwise comparison between Arabidopsis and tomato published cell type clusters. (A) for Figure 2a; (B) for Figure 2c; (C) for Figure 2d.

**Table S4:** The log transformation of p-values for statistical test across 15 species cell type clusters for Figure 3a.

**Table S5:** Number of cells and scRNA-seq publications used in this study.

