## SupplementaryFigures for "Orthologous marker groups reveal broad cell identity conservation across plant single-cell transcriptomes"

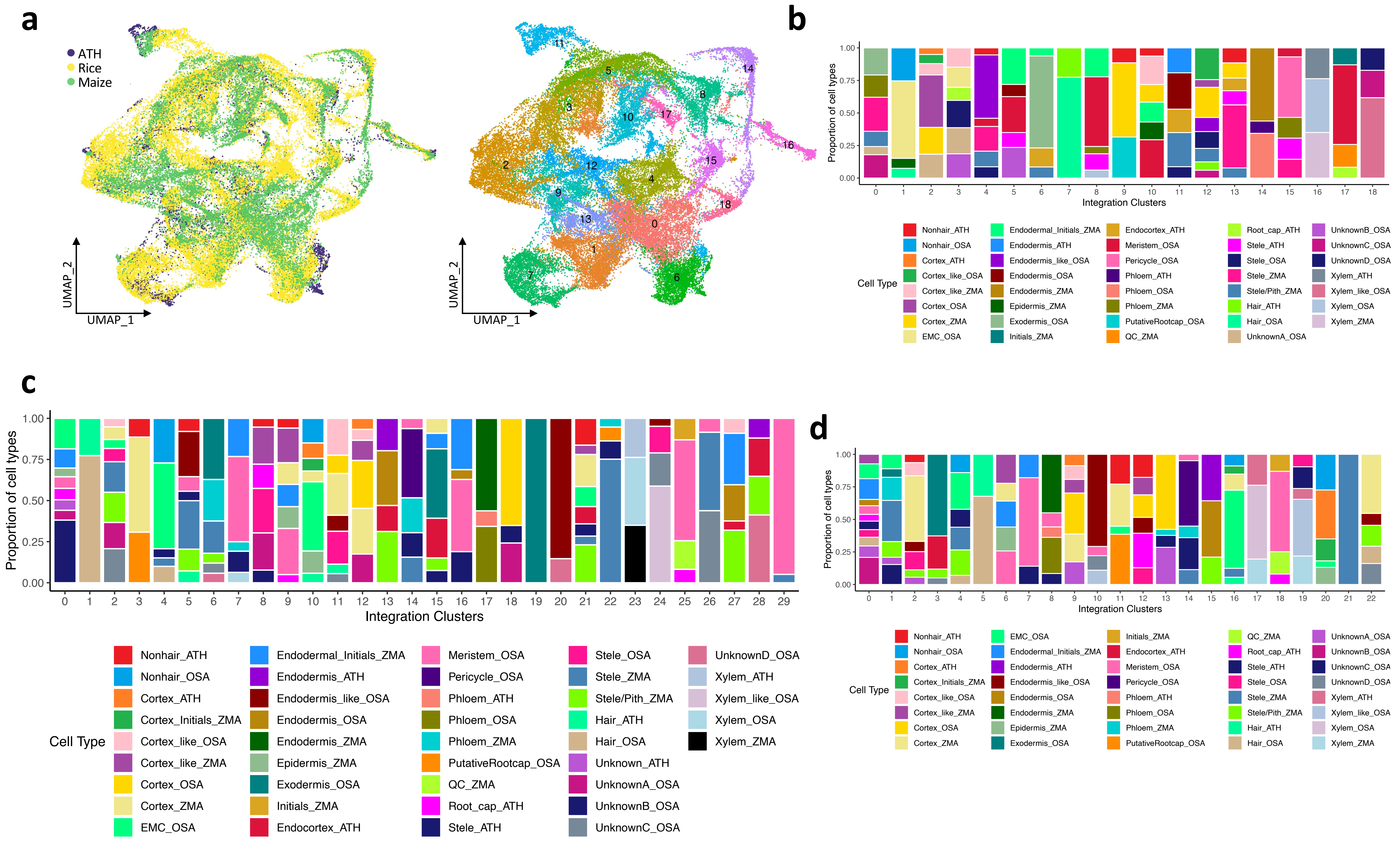


**Fig. S1:** **a**, integration of rice, maize, and Arabidopsis single-cell data. Left colors indicate different species. Right, colors indicate different clusters. **b**, bar plot shows all clusters are made of mixed cell types when using 19 clusters in the integrated map. Each color in the plot corresponds to a specific cell type cluster in a particular species, with the label indicating the cell type followed by the species ATH, OSA, or ZMA. **c**, Bar plot shows most clusters are made of mixed cell types when using 30 clusters. **d**, Bar plot shows most clusters are made of mixed cell types when applying Harmony integration.

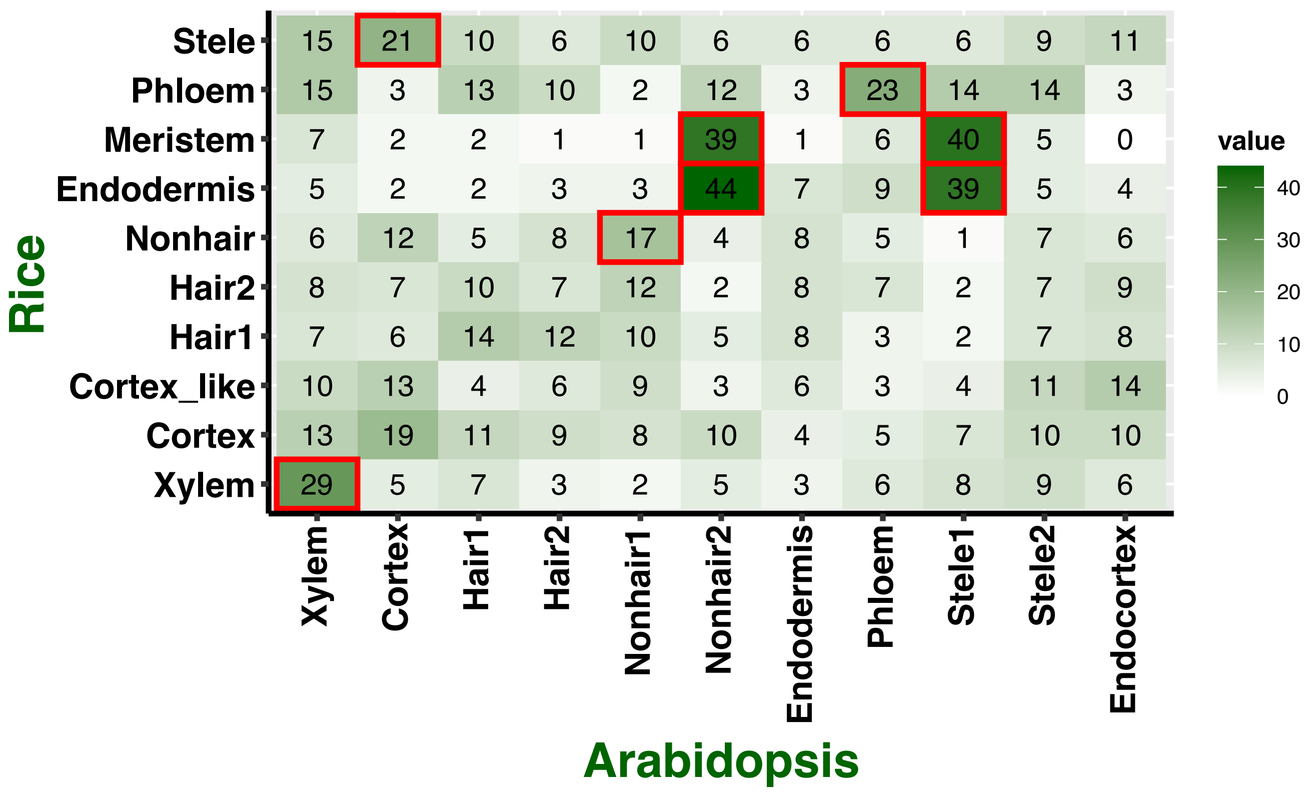


**Fig. S2:** Comparison of marker gene overlaps between Arabidopsis and rice using only 1-to-1 orthologous genes.

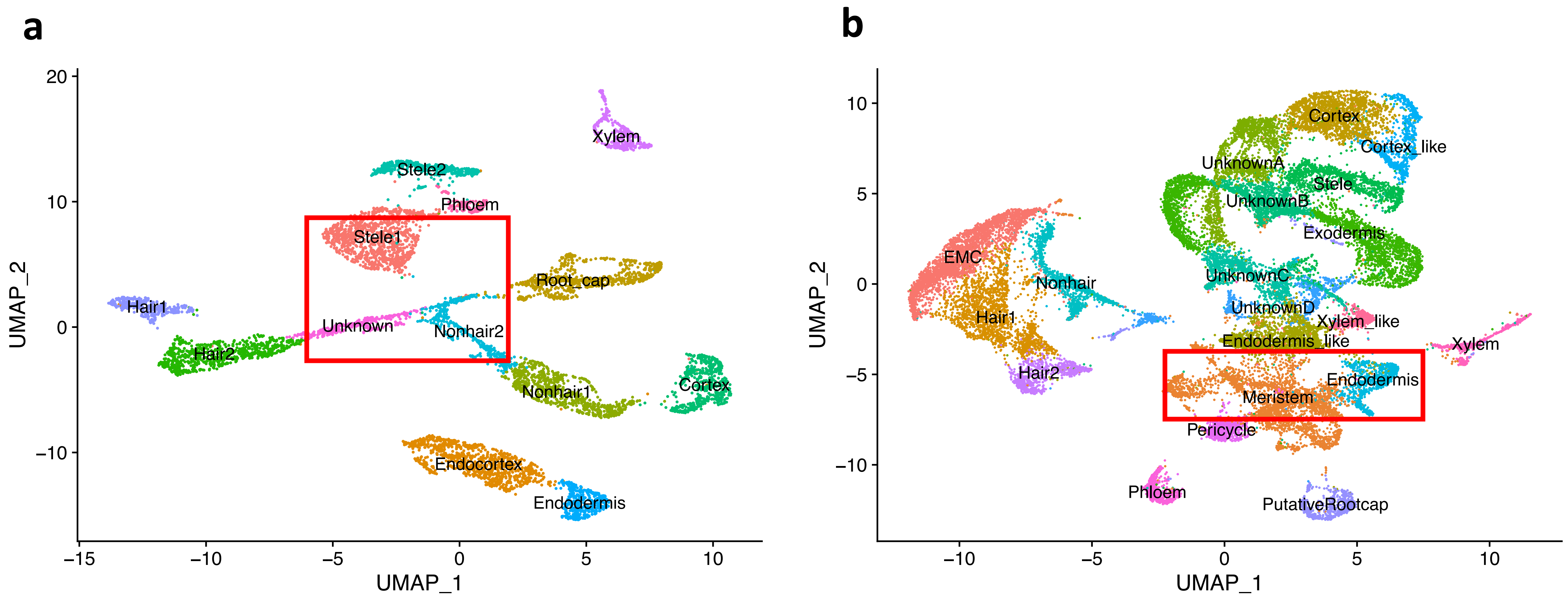


**Fig. S3:** The UMAP visualizations of the clustered cell groups of 7522 Arabidopsis cells **a**, and 27,426 Rice cells **b**, demonstrate that clusters such as the Stele1, Nonhair2, Meristem, and Endodermis, which are positioned at the center of the plots, exhibit a higher number of overlapping OMGs and this overlap can result in a mismatch between the cell type clusters of the two species (Fig. 1). Each dot represents an individual cell, with its color representing the respective cell type.

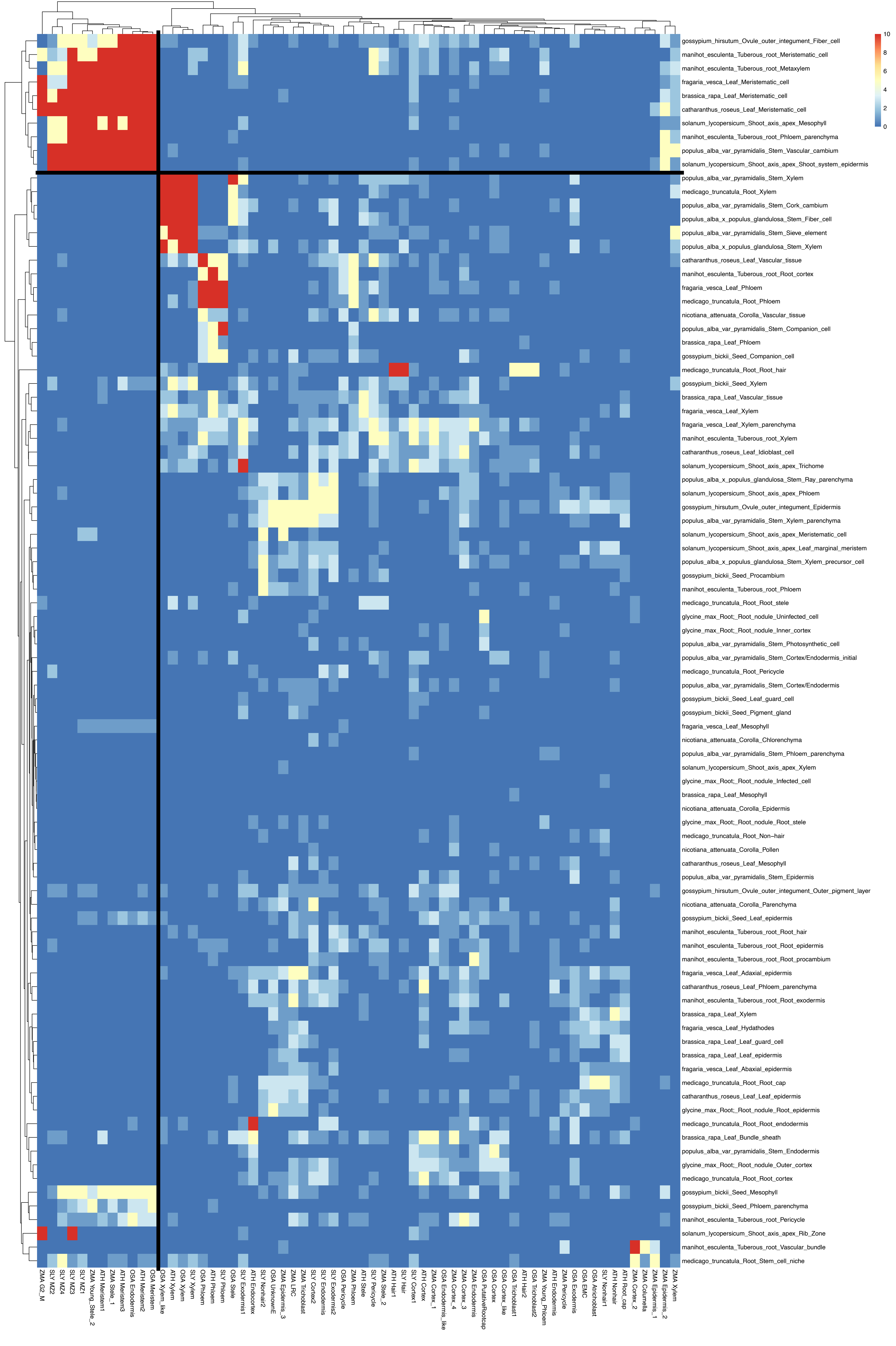


**Fig. S4:** cell type mapping heatmap between Arabidopsis, maize, rice, tomato (x axis), and 15 plant species (y axis). Red cluster on top left highlights cluster of cells from meristematic regions.

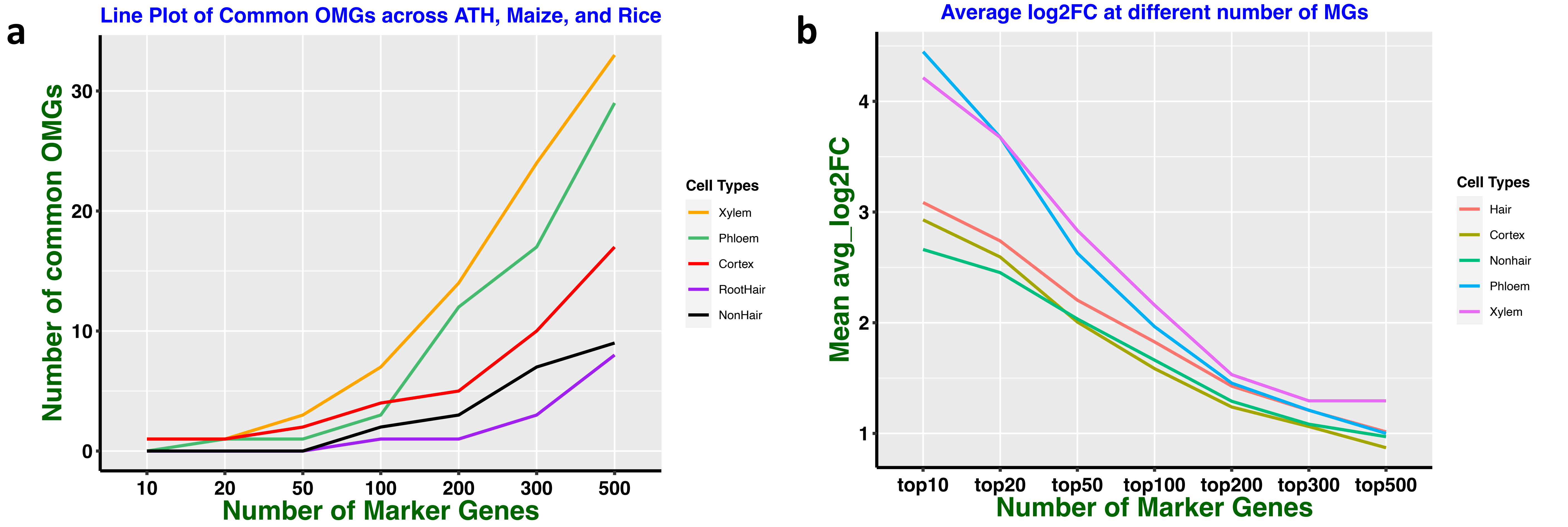


**Fig. S5: a,** The number of conserved orthologous marker genes (OMGs) across Arabidopsis, Maize, and Rice for varying numbers of marker genes. **b,** The average gene expression for varying number of top marker genes. These results showed that with increasing number of marker genes from each cluster, the number of OMGs also increase (a). However, the marker specificity, as measured by log2 fold change also decreases to 1 if we increase the marker number to 300 per cluster (b). If we decrease the marker number to 100 per cluster, the overlapping OMGs decreased to fewer than 10 for all cell types in this plot. These five cell types were selected because these are the only cell types that have conserved OMGs across three species: rice, maize and Arabidopsis.

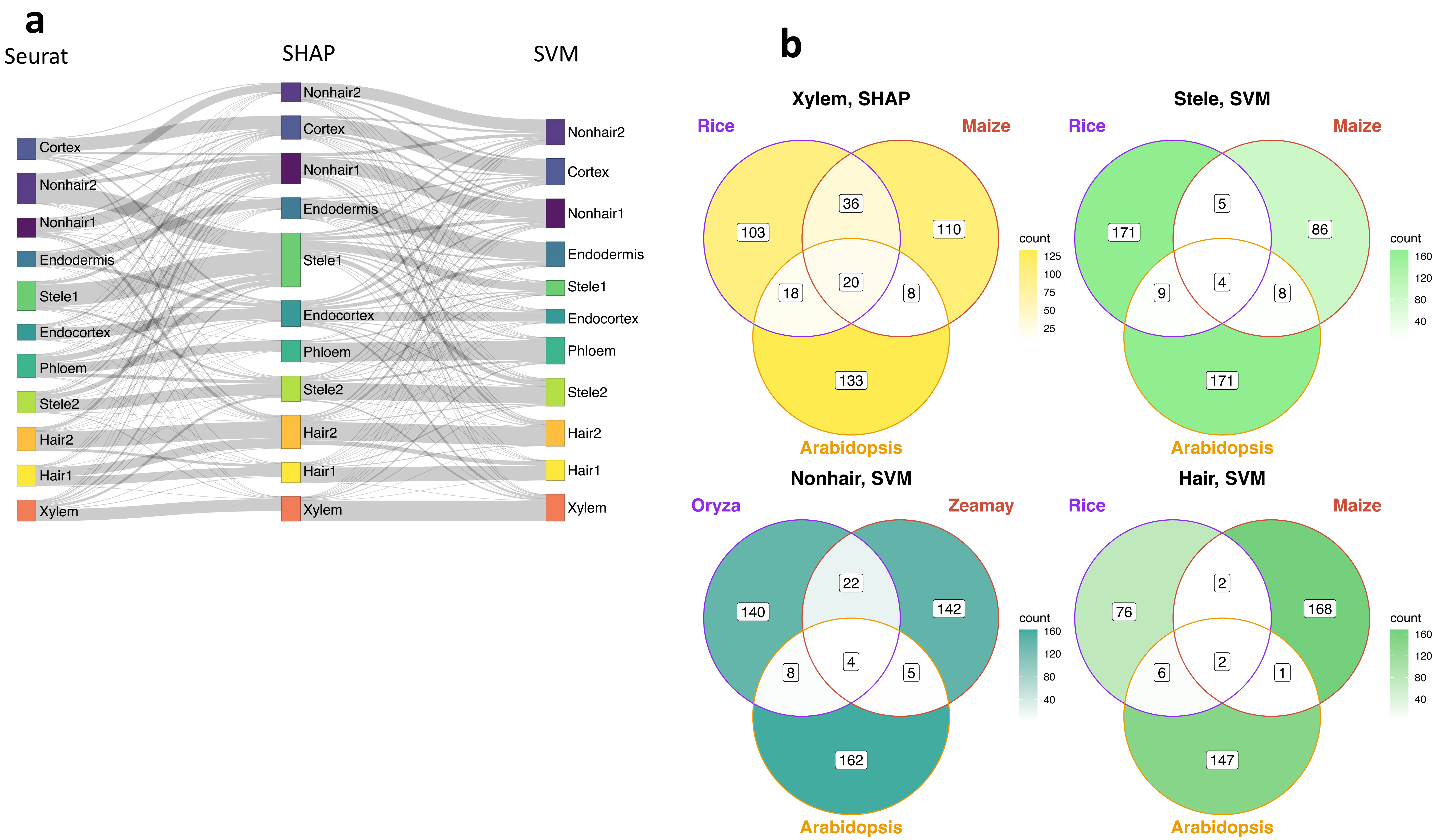


**Fig. S6: a**, comparison of the top 200 marker genes found by Seurat and machine learning methods. The height of the square and the thickness of the connecting lines represent the number of common marker genes found by the two methods, with each square's color representing a distinct cell type cluster. We found a high level of agreement between the ML-derived markers and fewer markers were shared between ML methods and the Seurat method. **b**, conserved OMGs found by machine learning approaches.

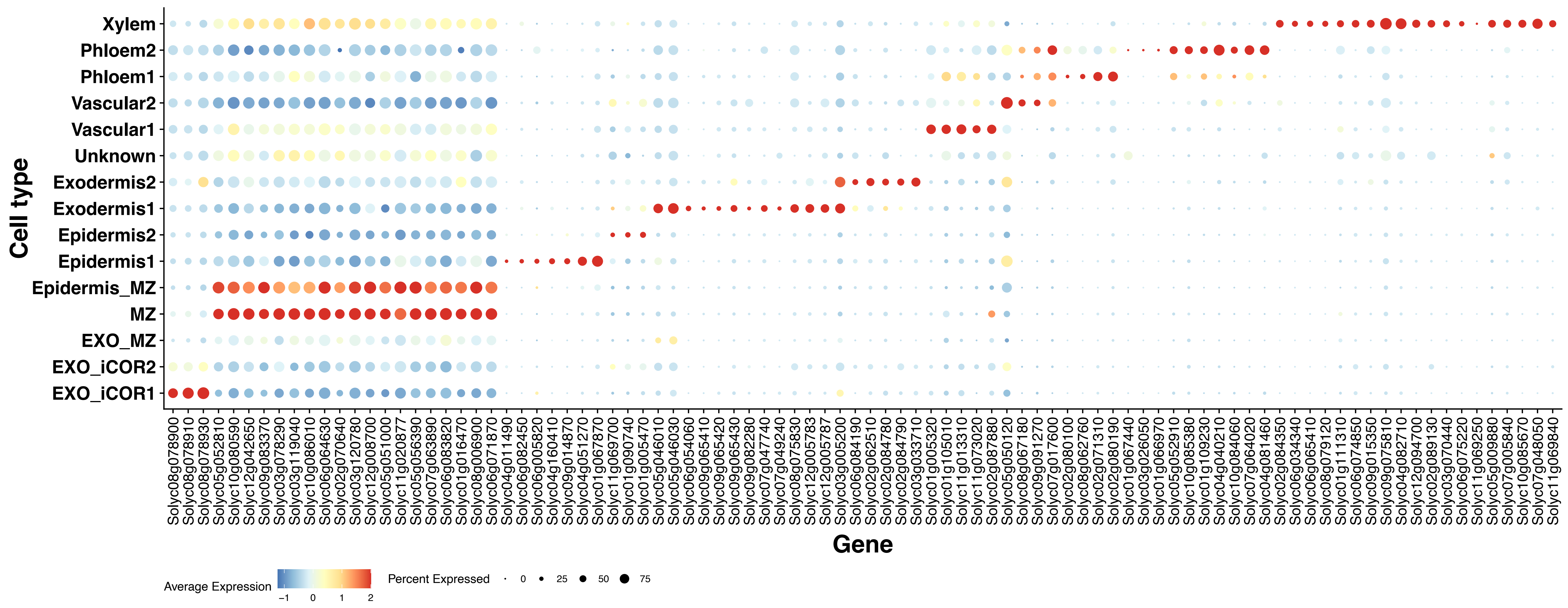


**Fig. S7:** Expression patterns of tomato cell-type marker genes which are derived from the conserved OMGs across species. The dot size represents the proportion of cells in a particular cluster expressing the marker gene, while the color of the dot indicates the level of expression. Red dots indicate high expression, while blue dots indicate low expression.

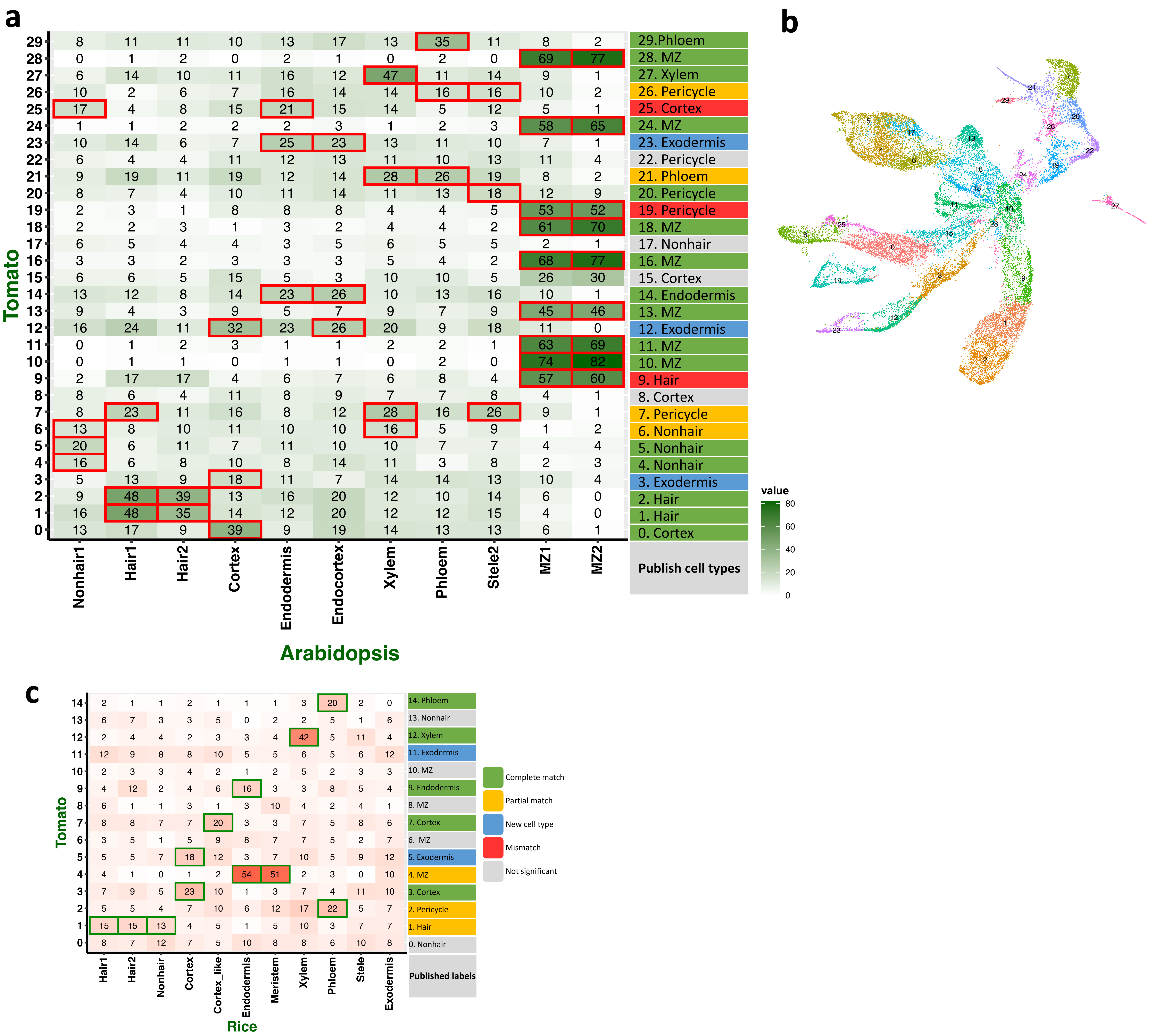


**Fig. S8: a**, using Arabidopsis OMGs to classify 30 tomato root cell types. **b**, UMAP of 30 tomato root cell cluters. **c**, Rice OMGs to predict 15 tomato root cell types.

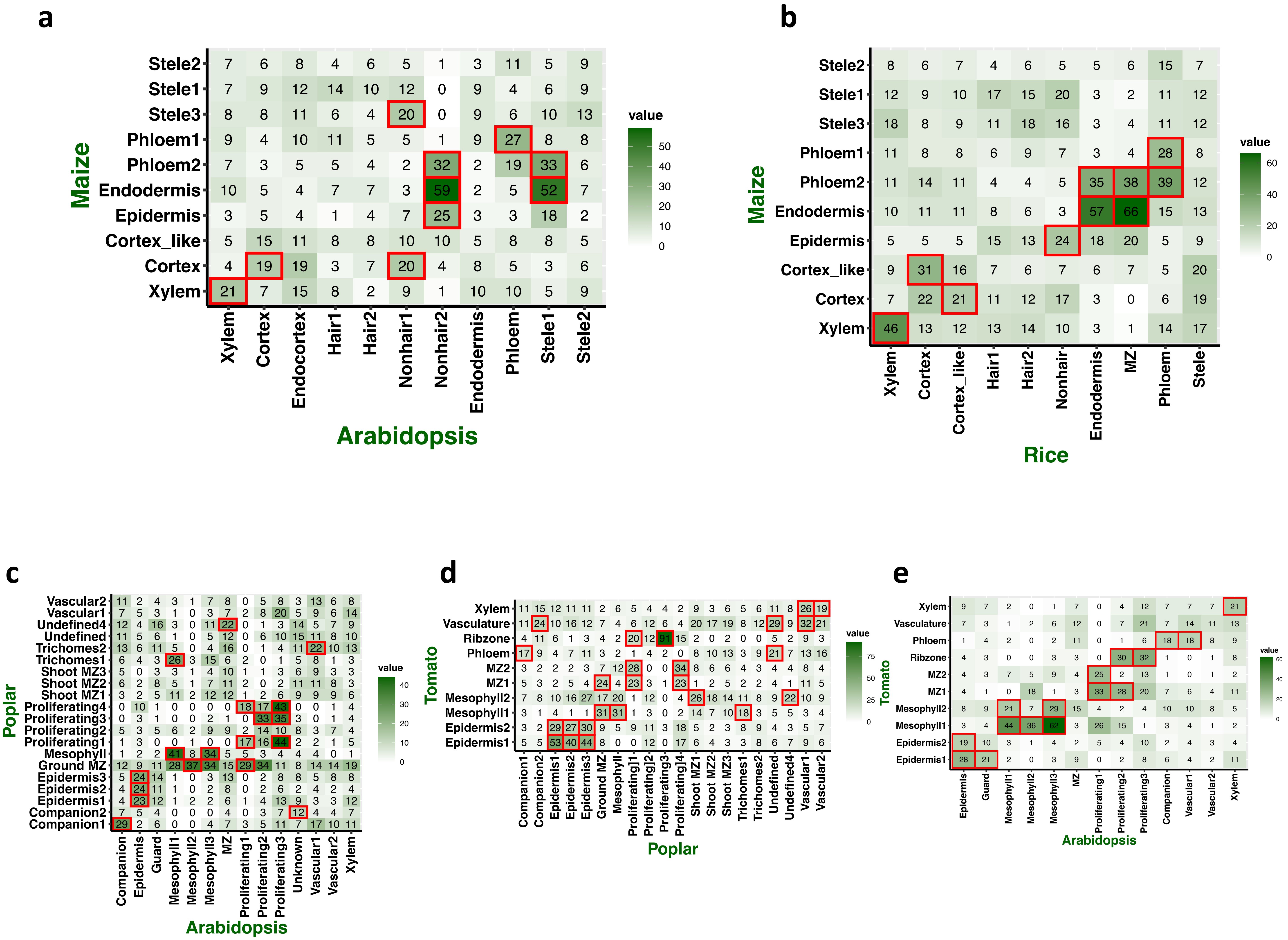


**Fig. S9:** Overlapping of OMGs between **a**, maize and Arabidopsis root, **b**, maize and rice root, **c**, Arabidopsis and poplar shoot, **d**, poplar and tomato shoot, and **e**, Arabidopsis and tomato shoot. The numbers displayed in each cell indicate the count of conserved marker genes between the two cell-type clusters. Significance is denoted by the red boxes.

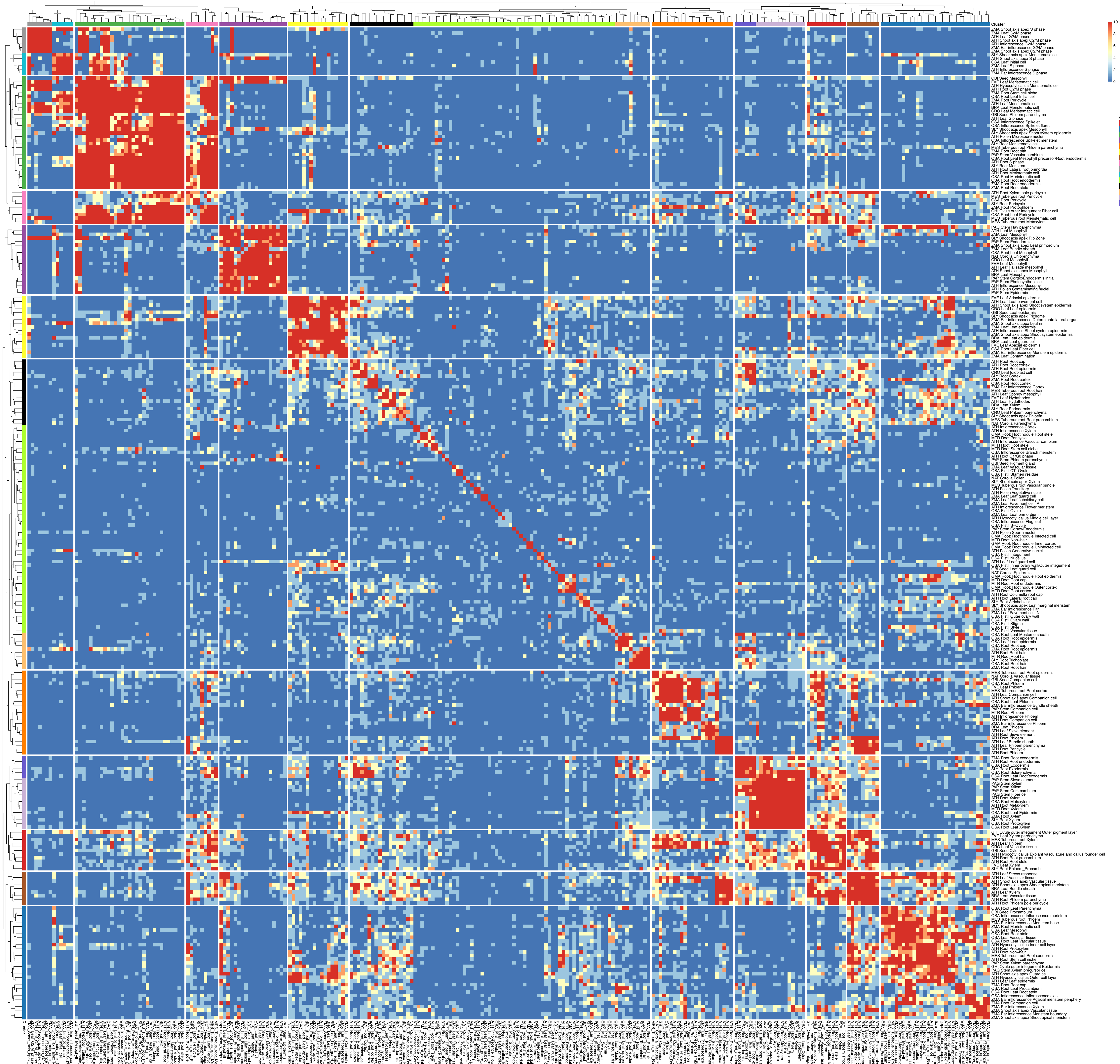

**Fig. S10:** cell type mapping across 15 diverse species based on the shared OMGs across species clusters with labels.

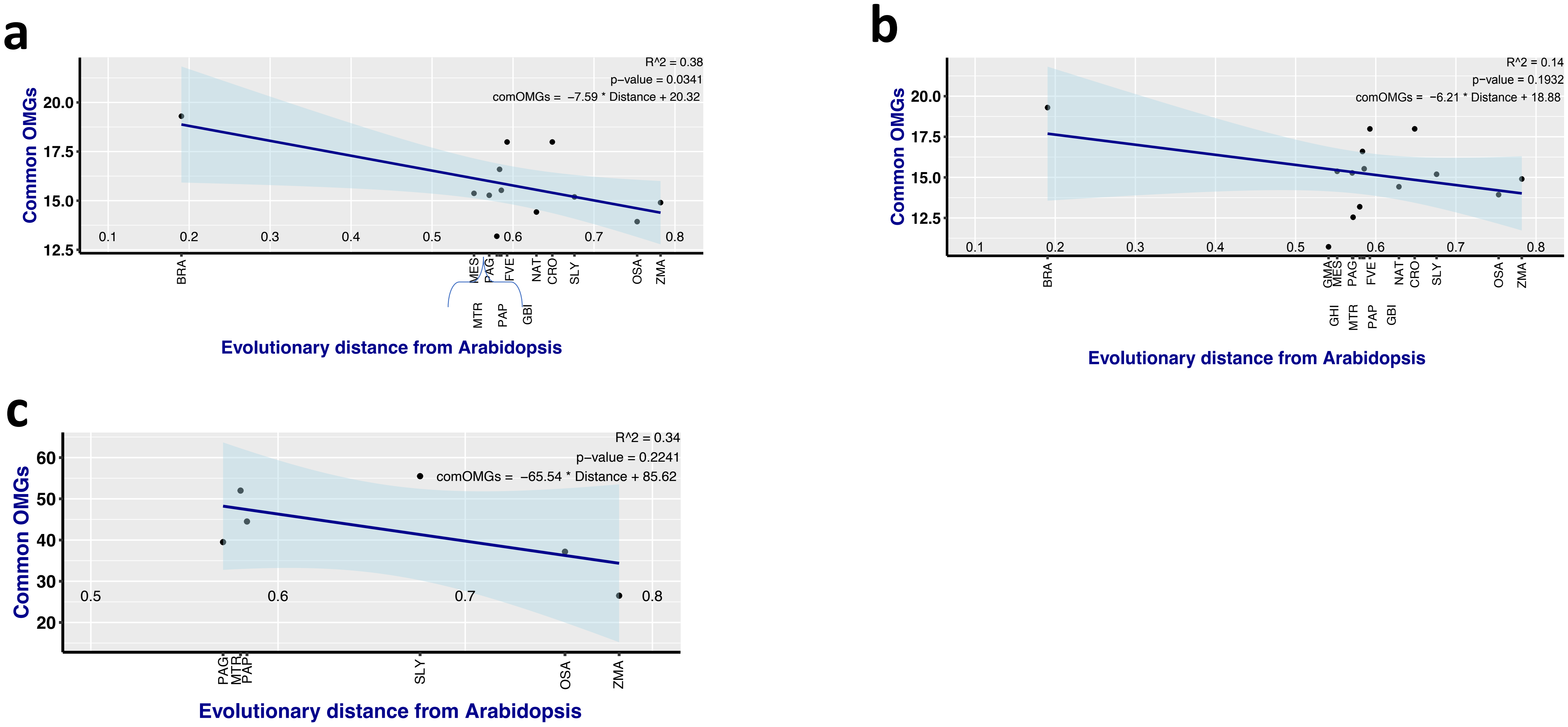


**Fig. S11:**

**a**, evolutionary distance vs. shared OMGs between 12 species and Arabidopsis. Soybean and upland cotton were excluded because their unique tissue types (root nodule, and ovule outer integument, respectively) and their limited data sets (six and three cell types, respectively) make them outliers. Excluding them from the analysis strengthened the relationship ($R^2$=0.38), further corroborated by a p-value <0.05.

**b**, evolutionary distance vs. shared OMGs between 14 species and Arabidopsis. Distant species generally share fewer OMGs than closer ones, though this correlation is not strong ($R^2$=0.14).

**c**, evolutionary distance vs. shared OMGs between root xylem in Arabidopsis and in other species. Xylem in root is the only cell type consistently observed across species in this dataset, displaying a correlation with evolutionary distance and shared OMGs, with a modest $R^2$ of 0.34.
